# How the Easter Egg Weevils Got Their Spots: Phylogenomics reveals Müllerian Mimicry in *Pachyrhynchus* (Coleoptera, Curculionidae)

**DOI:** 10.1101/2021.01.20.427190

**Authors:** Matthew H. Van Dam, Analyn Anzano Cabras, Athena W. Lam

**Affiliations:** Entomology Department, Institute for Biodiversity Science and Sustainability, California Academy of Sciences, 55 Music Concourse Dr., San Francisco, CA 94118, USA; Coleoptera Research Center, Institute for Biodiversity and Environment, University of Mindanao, Matina, Davao City, 8000, Philippines; Center for Comparative Genomics, Institute for Biodiversity Science and Sustainability, California Academy of Sciences, 55 Music Concourse Dr., San Francisco, CA 94118, USA

## Abstract

The evolutionary origins of mimicry in the Easter Egg weevil, *Pachyrhynchus,* have fascinated researchers since first noted more than a century ago by Alfred Russel Wallace. Müllerian mimicry, or mimicry in which two or more distasteful species look similar, is widespread throughout the animal kingdom. Given the varied but discrete color patterns in *Pachyrhynchus,* this genus presents one of the best opportunities to study the evolution of both perfect and imperfect mimicry. We analyzed more than 10,000 UCE loci using a novel partitioning strategy to resolve the relationships of closely related species in the genus. Our results indicate that many of the mimetic color patterns observed in sympatric species are due to convergent evolution. We suggest that this convergence is driven by frequency-dependent selection.

## INTRODUCTION

### Mimicry as a driver of species diversification

Mimicry is central to evolutionary biology and has important implications for various evolutionary processes, such as adaptive radiation^1^, coevolution^2^, niche partitioning^3,4^, population structuring^5^, adaptation of chemical defense^6^. In particular, Müllerian mimicry is hypothesized to be a primary driver of diversification resulting from mutualistic evolution^7–10^. This type of mimicry describes an antipredator strategy for groups of unpalatable species—by sharing similar color and pattern, each individual prey experiences a lower probability of being mistaken as a food source by predators^7^. Müllerian mimicry and aposematism have been the subject of many fascinating studies in a wide range of taxa: butterflies^11–13^, net-winged beetles^14^, velvet ants^15^, spiny plants^16^, dart frogs^17,18^, vipers^19^, coral snakes^20^, fish^21^, and possibly toxic birds^22^.

*Pachyrhynchus* is a diverse and charismatic group of beetles with elaborate patterning and iridescent colors (Figs. 1–2). Beetles in this genus have hard cuticles with fused elytra, which presents a line of defense against predation^23,24^, and they are hypothesized to be distasteful to their predators, which consist of birds, lizards, and frogs^23,25^. Their hard cuticle is derived in part through the bacteria endosymbiont *Nardonella,* which produces all precursors to tyrosine, a key amino acid for cuticular hardening^26^. *Pachyrhynchus* distinctive color patterns are structural colors produced by their scales’ inner nanostructure which scatters incident light^27^.

**Figure 1.**
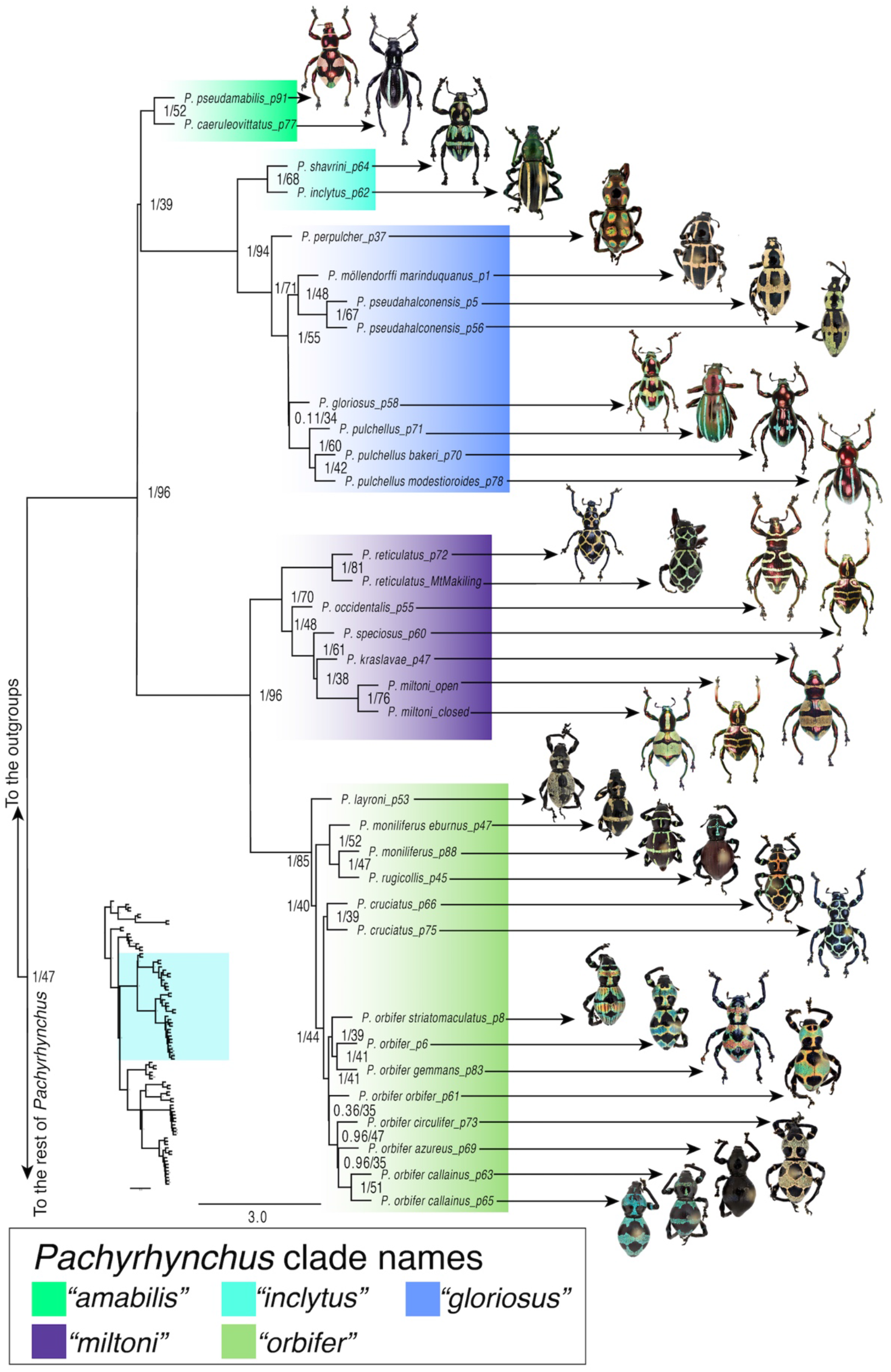
Phylogeny of the *Pachyrhynchus* ASTRAL species tree, part one. The other half of the tree is found in Fig. 2. Node labels correspond to local posterior probability (LPP) and normalized quartet support (NQS).

**Figure 2.**
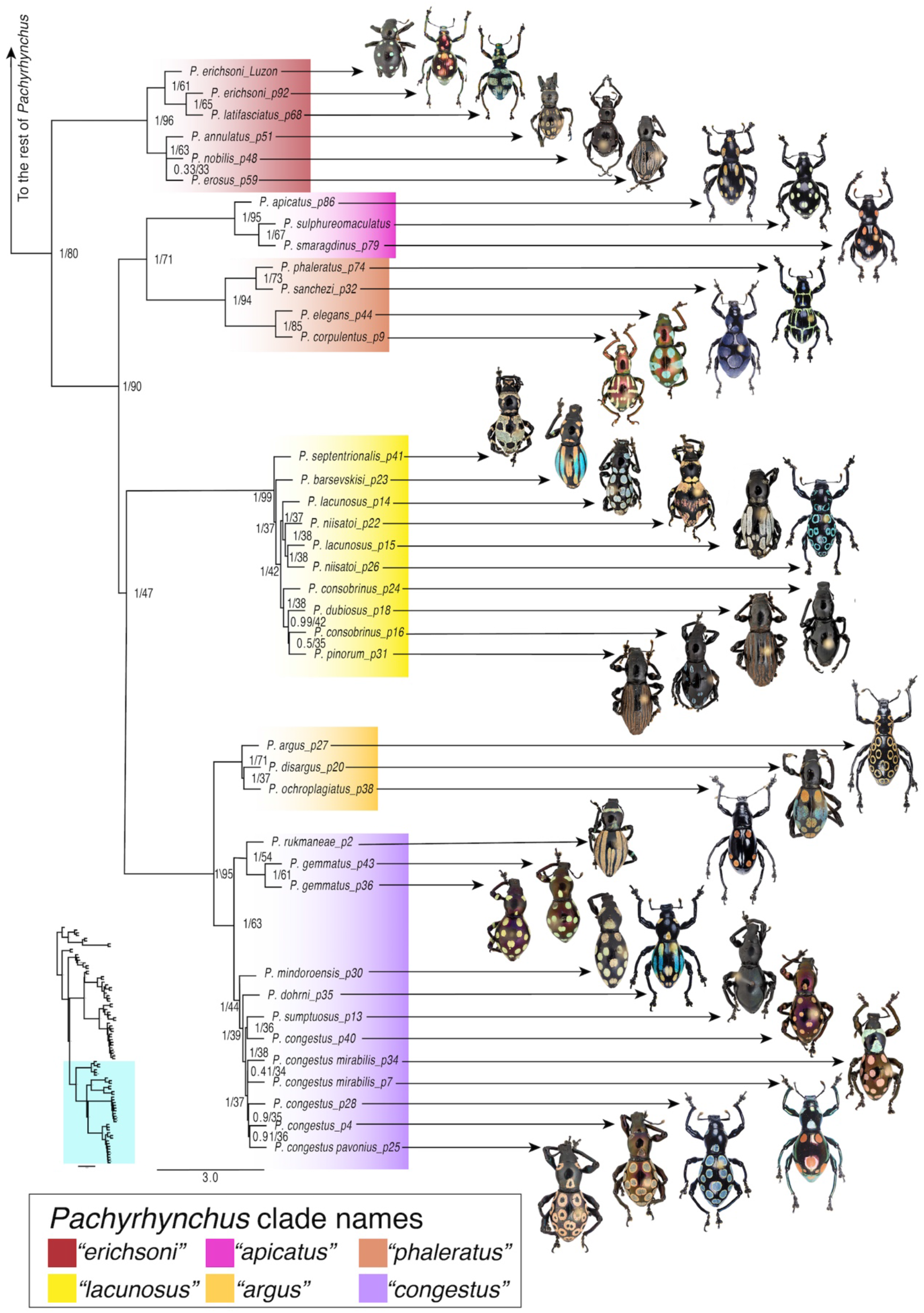
Phylogeny of the *Pachyrhynchus* ASTRAL species tree, part two. The other half of the tree is found in Fig. 1. Node labels correspond to local posterior probability (LPP) and normalized quartet support (NQS).

Both Batesian and Müllerian mimicry are associated with and found within *Pachyrhynchus.* The first record of mimicry among Pachyrhynchini was noted by Wallace^28^ when he observed sympatric species with the same colors and elytral patterns. It was also noted by Schultze where he provided a list of 19 sympatric species of *Pachyrhynchus, Metapocyrtus* (also in the Pachyrhynchini), and *Doliops* (Cerambycidae), all sharing the same coloration and patterns between genera^25,29^. He reported 14 additional sympatric species of *Metapocyrtus* exhibiting very similar elytral patterns, only distinguishable from each other by close inspection of diagnostic characters of the rostrum^29^. Their distinctive patterns are also observed in other unrelated weevils (e.g., *Polycatus, Eupyrgops, Neopyrgops, Alcidodes, Coptorhynchus, Calidiopsis).* Following the typical Batesian model, long-horned beetles (e.g., *Doliops, Paradoliops)* and even a cricket mimic *Pachyrhynchus’* aposematic signals are thought to aid in avoiding predation^28^. In this study, we explore patterns of Müllerian mimicry in *Pachyrhynchus* through detailed study of the group’s phylogenetics, biogeography, and ancestral state reconstructions.

### Pachyrhynchus’ natural history and taxonomy

The genus *Pachyrhynchus* was established based on the species *Pachyrhynchus moniliferus* from Luzon Island, Philippines, and placed in the tribe Pachyrhynchini (Germar). The tribe is entirely flightless with 18 described genera and more than 500 described species. *Pachyrhynchus* is endemic to oceanic islands but the entire genus is flightless. It is distributed in the Philippines (excluding Palawan), Ryukyu Island, Green and Orchid Island in Taiwan, and Talaud and Moluccas Island in Indonesia^25,30,31^, with the highest diversity in the Philippines. There are approximately 145 species of *Pachyrhynchus,* 93% of which are Philippine endemics^25,32^. Currently, despite the recent interest in the group^31–35^, *Pachyrhyhnchus* taxonomy and the relationships between different species remains poorly understood. Although naturalists and scientists have been collecting and describing *Pachyrhynchus* for centuries, there remains no concerted effort to study their phylogenetic relationships in a robust manner. While some species have been well quantified using morphometrics combined with DNA characters, the taxonomic scope of these studies are limited to only a few species found on Green and Orchid Islands of Taiwan^33^. The foundations of *Pachyrhynchus* taxonomy remains problematic, Heller^36^ and Schultze^25^ tried grouping species of *Pachyrhynchus* according to elytral color patterns. This was flawed because a heterogeneous set of unrelated species may lead to polyphyletic classifications. For example, classifications based on color pattern alone may cause over splitting of species because of intraspecific variation, or, conversely, over lumping of species because of convergent coloration and patterns. *Pachyrhynchus* species concepts remain largely untested by more complete morphological data and genetic data as well. We hope to provide a basic structure for the relationships within the genus to advance towards a more meaningful discussion on this group’s mimicry and biology.

### Biogeographic model

The Pleistocene Aggregate Island Complex (PAIC) which has been used as a biogeographic framework in the Philippines for decades, refers groups of islands in the Philippines with less than −120 m isobath separation that coalesced in the Pleistocene Epoch due to lowering of sea levels during glacial cycles^37–42^. These groups of islands are centers of biological endemism and share a high percentage of common floral and faunal elements. There are nine biogeographic subregions in the Philippines such as Batanes PAIC, Babuyanes PAIC, Luzon PAIC, Mindoro PAIC, Romblon PAIC, Palawan PAIC, Mindanao PAIC and Sulu PAIC^40^ of which the three largest biogeographically significant sub-provinces are Greater Luzon PAIC, Mindanao PAIC, and Greater Negros-Panay^38^. Studies on the distribution of mammals, birds, and herpetofauna have shown the consistency of PAIC boundaries on species distribution^38,40^. The connectedness of these islands allowed for the exchange of flora and fauna amongst the islands. The Philippines which has a unique geologic history including; paleo-island accretion, late Pleistocene sea-level fluctuations, and landmass connectivity has led to the isolation of lineages and restricted distributional patterns such as in the case of *Pachyrhynchus*.

Here, we use both dry pinned and newly collected specimens from the last 30 years and more than 10,000 UCE loci to gain a more thorough understanding of the *Pachyrhynchus* mimetic system. Our results are the first well-supported phylogeny of this charismatic and threatened beetle genus.

We explore the following study questions:

1. Müllerian Mimicry – We aim to identify how sympatric *Pachyrhynchus* species with similar patterns acquired their coloration. We tested if their similarity is due to inheritance alone or is the color pattern independently evolved in one or more sympatric lineages.
2. Intraspecific polymorphism – Some species of *Pachyrhynchus* display a striking polymorphism where some individuals possess solid maculations while others exhibit the same pattern but is not “filled” (i.e. same outline but lacking scales in the middle). We tested whether this trait is constrained to a single lineage within *Pachyrhynchus* or if it is more widespread throughout the genus.
3. Biogeography – Does *Pachyrhynchus* follow the Pleistocene Aggregate Island Complex (PAIC) hypothesis^37,43,44^, where speciation follows the interglacial periods when the land masses of the Philippines were at their most isolate in the last ~3MYA?

## METHODS

### DNA isolation and Quality Check

We used 71 pinned and 16 ethanol preserved specimens for this study. For a complete list of voucher specimens, see Supplemental Table S1. Voucher specimens are stored in two museum collections: the Coleoptera Research Center, University of Mindanao, Philippines (CRC) and the California Academy of Sciences, Entomology Department, USA (CASENT). To minimize contaminants, specimens were carefully dissected, excluding exoskeleton and beetle guts. In some specimens, contaminants are apparent upon examination (e.g., yeast, fungus and mites are observed in the body cavity), in those cases legs were removed and punctured to allow enzymatic (proteinase-K) digestion of soft tissue. DNA was extracted from the resulting tissues using the QIAamp micro kit (Qiagen, Germany) following the manufacturer’s protocol. We assessed the quantity of all isolated DNA using a Qubit 2.0 Fluorometer (Invitrogen, USA DNA quality (i.e., fragment size distributions) was determined using 1% agarose gel in newly collected specimens and historic samples with low starting concentrations, with a 2100 BioAnalyzer (Agilent Technologies, USA). We found that our starting DNA quality and quantity varied significantly among specimens: from 4.8 ng–5000 ng and 200 bp–50 kbp. This is not surprising as collection dates and storage conditions of each specimen varied widely.

High molecular weight (HMW) DNA for 10X Genomics library construction was extracted from a single newly collected, dry frozen, *P. miltoni* specimen, using MagAttract HMW DNA Kit (Qiagen, Germany) and following manufacturer’s protocol. DNA fragment size was quantified by pulsed-field capillary electrophoresis using the Femto Pulse System (Agilent Technologies, USA).

### Library Construction

We obtained paired-end Illumina data from 85 whole genome libraries, and an additional three from another studies (Van Dam et al. 2021, in prep). When necessary, DNA was sheared using a Covaris M220 (Covaris Inc., USA). Libraries were constructed with the NEBNext^®^ Ultra™II DNA Library Preparation kit (New England Biolabs Inc, USA) following the manufacturer’s protocol. To minimize PCR replicates in the final dataset, titration of the number of PCR cycles was performed as described in Belton et al. 2012^45^. Average sizes of the final libraries were around 250 bp (for libraries constructed from historic samples) to 400 bp (for libraries constructed using newly collected material).

The 10X Genomics linked-read library for Pachyrhynchus miltoni was prepared at QB3 Genomics at the University of California, Berkeley

### Low Coverage Genome Assembly

First, we trimmed adapters and low quality bases from the ends of our reads with *fastp* version-0.20.0 using the “detect_adapter_for_pe” setting^46^. Because we used two lanes of sequencing for some of our libraries, we concatenated the forward and reverse reads of these two lanes. We then concatenated unpaired reads into a single file for each species. Next, we used *SPAdes-3.11.1*^47^ to assemble the reads into scaffolds with k-mer values of 21, 33, 55, 77, 99 and 127 in the *SPAdes* assembly pipeline, using default settings for everything other than the memory (“-m 800”) and cpu threads (“-t 32”).

### Genome Assembly of Pachyrhynchus miltoni

We used 594M illumina 2×150 paired reads which had been barcoded by the 10X Genomics Chromium instrument. The 10X Genomics linked-read assembly was constructed with Supernova v2.0.1^48^ with default settings.

### Ultraconserved Element (UCE) Marker Design

We designed a custom ultraconserved element (UCE) probe set using the PHYLUCE pipeline^49,50^. To maximize the effectiveness of the probes, we selected individuals that spanned the phylogenetic diversity of the Pachyrhynchini. For the base taxon, we used the chromosome level genome assembly of *Pachyrhynchus sulphureomaculatus* Schultze, 1922^52^. This taxon was used to select the initial bates because: 1) preliminary investigation of *P. sulphureomaculatus* indicated that it is not recovered at distal portions of the *Pachyrhynchus* phylogeny, which has been demonstrated to increase the number of UCE loci captured^52^, 2) the genome is complete and soft masked for repetitive elements, and 3) perhaps most important, the genome is free of contamination, which can lead to off-target loci capture (Van Dam et al. 2021, in prep). Soft masked genome is critical because the PHYLUCE pipeline only screens for repetitive DNA if the base genome is soft masked. If the genome is not soft masked, probes may be designed from paralogous loci. To diminish, as much as possible, the possibility of designing loci from non-target species in the probes themselves, we used a stringent screening method for our probes, selecting only loci that were recovered in our base taxon (Van Dam et al. 2021, in prep). We included eight *Pachyrhynchus* species from different species groups (Supplemental Figure S1). We selected two outgroup species, close relatives to *Pachyrhynchus*: *Coptorhynchus* and *Oribius* (Celeuthetini)^53^, to ensure that the probe set is relatively universal across the phylogenetic diversity of our target species, for a total dataset of ten species.

### Extracting UCE Loci and Alignment Construction

To match and extract our probe set from the *Pachyrhynchus* scaffolds and then align our loci we followed the PHYLUCE pipeline^49^ using default settings unless otherwise noted. We used the PHYLUCE script “phyluce_probe_run_multiple_lastzs_sqlite” with an “identity” of 60. After matching probes to scaffolds, we used “phyluce_probe_slice_sequence_from_genomes” to extract the flanking 500 bases around our probes. After the initial alignment step using *mafft*^54^ we used “phyluce_align_get_trimal_trimmed_alignments_from_untrimmed” to internally trim our matrices. This step uses *trimAI*^55^ to help trim ambiguously aligned sites in the alignments. We used “phyluce_align_get_only_loci_with_min_taxa” to select loci with a minimum of 50% complete matrices. Lastly, we used “phyluce_align_format_nexus_files_for_raxm” to produce the final concatenated matrix.

### UCE Phylogenomics

We used two different types of analyses for phylogenetic reconstruction: (1) a concatenated analysis using RAxML-NG v1.0.0^56^, and (2) a summary species tree analyses using ASTRAL-MP v5.7.4^57,58^.

### Concatenated Phylogenetic Analyses

We used the General Time Reversible + gamma (GTRGAMMA) site rate substitution model across our alignment. We used 10 independent parsimony-based starting trees for our maximum-likelihood (ML) searches in RAxML-NG. Non-parametric bootstrap replicates (BS) were done using the autoMRE option, with a maximum of 200 replicates to optimize the number of bootstrap replicates for this large dataset. Lastly, we mapped the bootstrap replicate values onto the best-scoring ML tree.

### Species Tree Analyses

#### Identifying UCE loci within genomic windows

We used methodologies that are similar to Van Dam et al. 2020^51^ for combining UCE loci. First, we identified UCE loci that are within 25 kb non-overlapping windows, as similar window sizes were used in Edelman et al. 2019^59^. To accomplish this, we divided up the base genome’s chromosomes into 25 kb non-overlapping windows using the *“tileGenome”* function in the *GenomicRanges* package^60^ in *R*. Next, we extracted the genomic coordinates from these windows and the coordinates produced in PHYLUCE for the probe set, and then we identified overlapping ranges using the *“GRanges”* function in the *GenomicRanges* package. If there was a UCE that overlapped two windows, it was combined with the adjoining UCEs to maximize the number of UCEs within a window.

#### Partitioning procedures

Data preperation before partitioning and partitioning procedures were carried out following Van Dam et al. 2017^62^. Before we partitioned UCE loci, we first used the R package *ips*^61^ with an *R* script from Van Dam et al. 2017^62^. We removed any columns composed exclusively of “-”,”n” and/or “?” using the “deleteEmptyCells” function, followed by removing any ragged ends of the matrix with the *“trimEnds”* function, with a minimum of four taxa present in the alignment. Below we expand on partitioning procedures to consider the potential site rate heterogeneity found in UCE loci as well as partition loci found in the same genomic window or potentially the same gene^51^. We use an alternative partitioning scheme from Van Dam et al. 2017 and Tagliacollo and Lanfear 2018^62,63^. Although the central core regions of UCEs tend to be more conserved than the flanking regions^49,62–64^, this observed amount of variability is greatly reduced due to internal trimming procedures in the PHYLUCE pipeline, which is necessary to accurately align loci. Internal trimming reduces the variability of informative sites to present a relatively even distribution across these loci^64^, essentially taking the U-shaped distribution and reducing it to a more or less flat line^64^. With this in mind, we unlinked the character sets of the flanking regions and used *PartitionFinder2* v2.1.1^65^ to group these separate partitions based on the best fitting model and substitution rates for the data. First, we divided the loci into a central core region of 80 bp because some loci are reduced by internal trimming to be less than the original 160 bp of the probes. Next, we divided each of the flanking regions into 5 separate character sets based on their proportion of sequence length (5 to the left of the central core and 5 to the right). For the loci that shared genomic windows, we combined character sets; thus, if two loci were concatenated, there would be 22-character sets total. These character sets were then input into *PartitionFinder2* for partitioning and model selection. Because RAxML-NG uses a more diverse set of nucleotide substitution models than previous versions of RAxML^56^, we tested for the best model fit from a total of 39 different models (Supplemental material, model list).

#### Gene tree and species tree reconstruction

We used ten independent parsimony-based starting trees for our maximum likelihood (ML) searches in RAxML-NG. We then performed 100 non-parametric bootstrap replicates, followed by mapping the bootstrap replicate values onto the best-scoring ML tree. Next, we collapsed/contracted branches in the gene trees with BS ≤20 using newickutils^66^; which has been demonstrated to have a strong positive impact on the accuracy of species tree reconstruction^67,68^. These resulting trees were used in species tree reconstruction. We used ASTRAL-MP with the default settings to reconstruct the species tree and annotate the tree with support values calculated for the normalized quartet support (NQS) and local posterior probability (LPP)^69^.

### Divergence Dating Biogeography

We used MCMCTREE^70^ to perform our divergence dating analyses. We used the topology from our ASTRAL analyses as the starting topology for the MCMCTREE. No fossils exist for our ingroup or near relatives, so we used a geological calibration for the maximum age of the Philippine Islands. We used 25–30 Ma as the root node of the Pachyrhynchini, an approximate date for the emergence of the Philippine proto-islands proposed by Hall 2002. We used MCMCtreeR_71_ to estimate a normal distribution around the maximum age of our calibration point as well as to format the tree file for MCMCTREE. Next, we used the aforementioned tree to obtain a rough estimate of the substitution rate using basml^72^. To help accomplish this, we randomly selected 300 loci, using many more will prevent the analysis from completing, as well as using more data is not necessary to approximate the uncertainty of the divergence dates^72,73^. Finally, we estimated the gradient and Hessian of the branch lengths^74^ to assist in the final estimation of our divergence dates.

To reconstruct the broad scale biogeographical patterns of *Pachyrhynchus,* we used BioGeoBEARS v1.1.2^75^. Because biogeographic model selection is sensitive to duplicate taxa, we removed all potential duplicates from the same metapopulation lineage/species^75^. We wanted to examine if the biogeography followed the Pleistocene Aggregate Island Complex (PAIC) hypothesis^37,43,44^, and thus we defined our areas according to the PAIC scheme. Many of the defined areas are congruent with the various geological histories of the major island groups^42,76–78^ (Fig. 4). We followed this scheme except for the island of Marinduque because it contains a number of unique species which do not have any obvious close relatives in the Luzon PAIC. For this island, we were curious to see its colonization history separate from the Luzon PAIC. We initially examined three different biogeographic models, DEC^79^, DIVA-like^80^, BAYAREA-like^81^ (see Table 3), and also included the “+J” parameter for founder-event/jump speciation at cladogenesis events^75^. This parameter has been demonstrated to greatly improve model fit for island- and island-like systems^75,82,83^. We used the Akaike information criterion corrected for sample size (AICc) to identify which model best explained our data given the number of free parameters.

**Table 1.** Count of UCEs by the number of UCEs concatenated in a “set”. Upper row gives the number of UCEs concatenated in a 25 kb non-overlapping window, and the lower row is the count by category.

|  |  |  |  |  |  |  |
| --- | --- | --- | --- | --- | --- | --- |
| Length of UCE sets | 2 | 3 | 4 | 5 | 6 | 7 |
| Number of sets by length | 1087 | 184 | 44 | 11 | 4 | 2 |

**Table 2.** Average bootstrap support by locus type. “Sets” are concatenated UCEs found in a 25kb non-overlapping sliding window bin, and “Singles” are those where only a single UCE occupies a 25kb bin in the *P. sulphureomaculatus* base genome.

| Loci type | ABS_mean | ABS_median | ABS_95%_CI | Number of loci |
| --- | --- | --- | --- | --- |
| Sets | 55.4 | 58 | 0.67796 | 1332 |
| Singles | 48.1 | 51.9 | 0.41227 | 7113 |

**Table 3.** BioGeoBEARS model results table.

| Biogeographic model | LnL | numparams | d | e | j | AICc | AICc_wt |
| --- | --- | --- | --- | --- | --- | --- | --- |
| DEC | -80.65 | 2 | 0.0069 | 1.00E-12 | 0 | 165.5 | 2.60E-10 |
| DEC+J | -58.12 | 3 | 1.00E-12 | 1.00E-12 | 0.038 | 122.7 | 0.53 |
| DIVALIKE | -74.52 | 2 | 0.0089 | 1.00E-12 | 0 | 153.2 | 1.20E-07 |
| DIVALIKE+J | -58.85 | 3 | 1.00E-12 | 1.00E-12 | 0.039 | 124.1 | 0.25 |
| BAYAREALIKE | -100.1 | 2 | 0.0058 | 0.058 | 0 | 204.5 | 9.00E-19 |
| BAYAREALIKE+J | -59.01 | 3 | 1.00E-07 | 1.00E-12 | 0.038 | 124.4 | 0.22 |

### Ancestral State Reconstruction of Color Patterns

To identify instances of mimicry and/or convergence in the color pattern of *Pachyrhynchus,* we used the R package *phytools* v0.7.70 “fitMk”^84^ to reconstruct ancestral character states. We categorized the color pattern into 11 different states (Fig. 5). We define 11 different color patterns on their elytra as follows: 1) “Black”, species whose integument is black or almost entirely black. 2) “Rainbow” elongate linear bands or elongate maculations that nearly touch composed of yellow/orange color at distal and apical ends gradually transitioning to blue in the center. 3) “Vertical bars” elongate linear bands or elongate maculations that nearly touch composed of a single color running the length of the elytra. 4) “Filled Moroccan tile”, large central patches of color with three apical and three distal areas not covered by colored scales. 5) “Open bands” three pairs of lines that circumscribe an ovoid shape or run the width of an elytra. 6) “Moroccan tile” net-like pattern of lines. 7) “Checker” two vertical lines near apex of elytra with a horizontal central line and two vertical lines near anterior end of elytra. 8) “Spots” at least three pairs of circular maculations. 9) “Irregular grid” two–four pairs of short convergent or parallel lines near apex, a central horizontal band, and two–four pairs of short convergent lines at the anterior end of the elytra. 10) “Vertical lines” narrow lines running the length of the elytra. 11) “Filled bands” three pairs of broad ovoid maculations running roughly the width of an elytra.

We then selected the best fitting model (equal rates ER, symmetric backward and forward rates SYM, or all-rates-different for transitions ARD) using the AIC weights as our selection criteria. All models treated character transitions as unordered. Next, we simulated 1000 different character histories using *phytools’* “make.simmap” function using the “mcmc” option to estimate the transition rate matrix, under our best fitting model. Lastly, because some *Pachyrhynchus* species possess discrete polymorphisms, we wanted to examine the history of this trait as well. The polymorphic trait in *Pachyrhynchus* presents itself as the color pattern being “filled” (solid center) or “open” (outlined) maculations (Fig. 4). The addition of this polymorphic state to the previous 11 character states (Fig. 4) resulted in a transition matrix that was much too large to analyze. To examine how widespread open or filled maculations are across the phylogeny, we coded it as two different polymorphic states (filled, not filled, or species that have both states). We performed the same model selection and ancestral state reconstruction methods as above but using *phytools’ “fitpolyMk”* and also included the model *“transient’,* where the rates of gaining or losing polymorphic states differ. We combined this information with their biogeography to identify whether color patterns were due to inheritance (related species look alike via shared ancestry of trait), convergence (species look alike but independently evolve a trait in allopatry), or mimetic convergence (species look alike, and of those that are similar, at least one lineage independently evolved the trait in sympatry).

**Figure 4.**
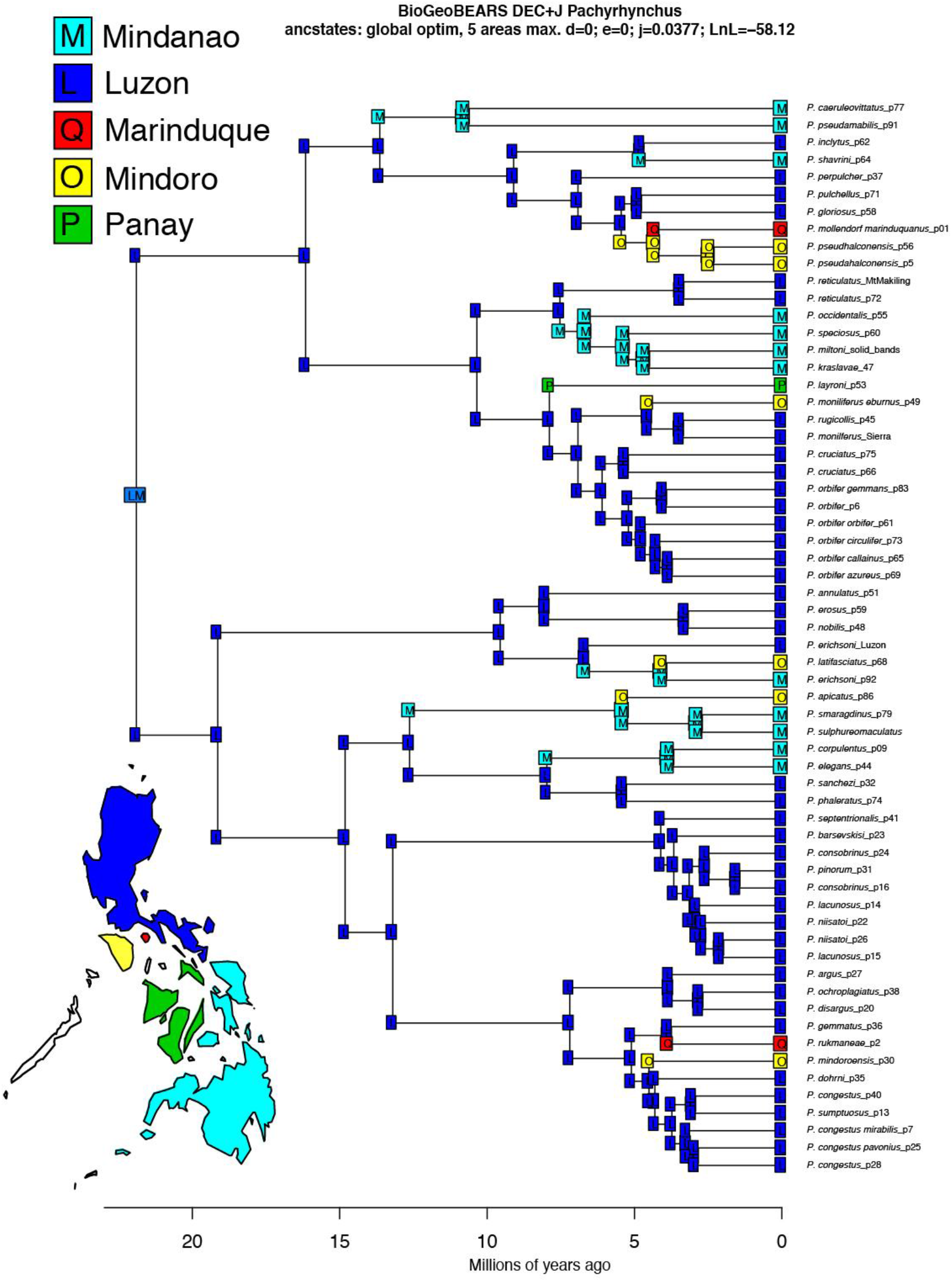
Biogeographic ancestral area reconstruction using *BioGeoBEARS “DEC+J”* model for *Pachyrhynchus.* Color codes correspond to Pleistocene Aggregate Island Complex (PAIC). Lower left color-coded map of the Philippines.

## RESULTS

### Genome Assembly of Pachyrhynchus miltoni

The 10X Genomics linked-read assembly resulted in a total length of 2,204,292,973 bp (roughly the same size as the *P. sulphureomaculatus* base genome^51^), with an N50 of 10,763 bp and 368,426 scaffolds. To use the assembly in PHYLUCE, we modified the headers of the scaffolds to resemble those from SPAdes, giving each a unique name and matching sequence length of the scaffold.

### Probe Design, Sequencing Results, and UCE Loci Recovery

Our probe design selected a total of 12,522 UCE loci shared by all 11 taxa. Our sequencing results were largely successful with only one DNA library producing less than a million reads, and this sample was removed from all subsequent analyses. Filtering loci that contain ≥ 50% complete matrices resulted in 10,108 of 12,323 total loci recovered. We sequenced an average of 77,990,397 ±11,627,118 95% CI, paired reads per sample (Supplementary Fig. S2), with a final 50% complete matrix with a mean of 8,756 ± 698 95% CI per sample (Supplementary Fig. S2). These statistics exclude loci recovered from the long read or pseudo long read assemblies, which recovered an average of ~11,000 UCEs (Supplementary Table S1). We found a total of 2,995 UCEs sets in 25 kb windows, with a total of 1,332 windows containing multiple UCEs and 7,113 UCEs not in sets (Supplementary Fig. S2).

### Phylogenetic Analyses

#### Concatenated analysis

Our alignment totaled 6,353,380 bp in length. The analysis completed 200 non-parametric bootstrap replicates, and we provided the phylogenetic hypothesis for the genus in Supplemental Fig 1S. BS support was relatively high throughout the tree (Supplementary Fig. S1), with strong support along the backbone (BS =100%). The genus *Pachyrhynchus* was recovered as monophyletic, but the genus *Metapocyrtus* was not. The genus *Pantorhytes* is sister to the Philippine members of the Pachyrhynchini, and the Celuthenini taxa are sister to Pachyrhynchini.

#### Species tree analyses

We present the first genus wide phylogeny, and recovered 11 major lineages (Figs. 1–2). We found surprising patterns that contradicts the idea that pattern alone is a sufficient character to group *Pachyrhynchus* species. For example, *P. speciosus,* previously described as a single species, was composed of several unrelated lineages^85^. In addition, *P. reticulatus* and *P. cruciatus* were actually rather distantly related despite similar “Moroccan tile” like patterns (Fig. 1).

The partitioning results of the gene trees showed that most UCEs require more than three partitions; with the mean number of partitions in single UCEs being 3.6 ± 0.04, and the mean number of partitions in UCE sets 17.8 ± 0.65. These results indicate that UCE loci have different rates and models of nucleotide substitutions as well, not just a central core and symmetrically variable flanking regions. In addition, the substitution models selected were varied, with the traditional GTR+G selected second-most frequently (Fig. 3). The topology of the species tree reconstructed with ASTRAL had an identical backbone to the tree from the concatenated analysis and only a few differences at nodes near the tips (Supplementary Fig. S3), but all species remained in the same clades (Figs. 1–2) in both trees. The local posterior probabilities (LPP)^69^ were similar to the BS values of the RAxML-concatenated analysis, with high support along the backbone of the tree and lower support near the tips, where conflict between topologies was typically due to short internode distances (Supplementary Fig. S3). Normalized quartet support (NQS) values were similar to the other support values (Figs. 1–2) with most of the gene trees congruent with the species tree topology along the backbone, as well as at nodes separated by relatively long internode distances. However, nodes separated by very short distances had more conflict among the gene tree topologies, with only ~35-40% of the gene tree quartets in agreement. The number of loci and our computer resources impaired the completion of the 100 BS replicates. However, the statistics we used are a more reliable measure of gene tree conflict and support for species trees^69^.

**Figure 3.**
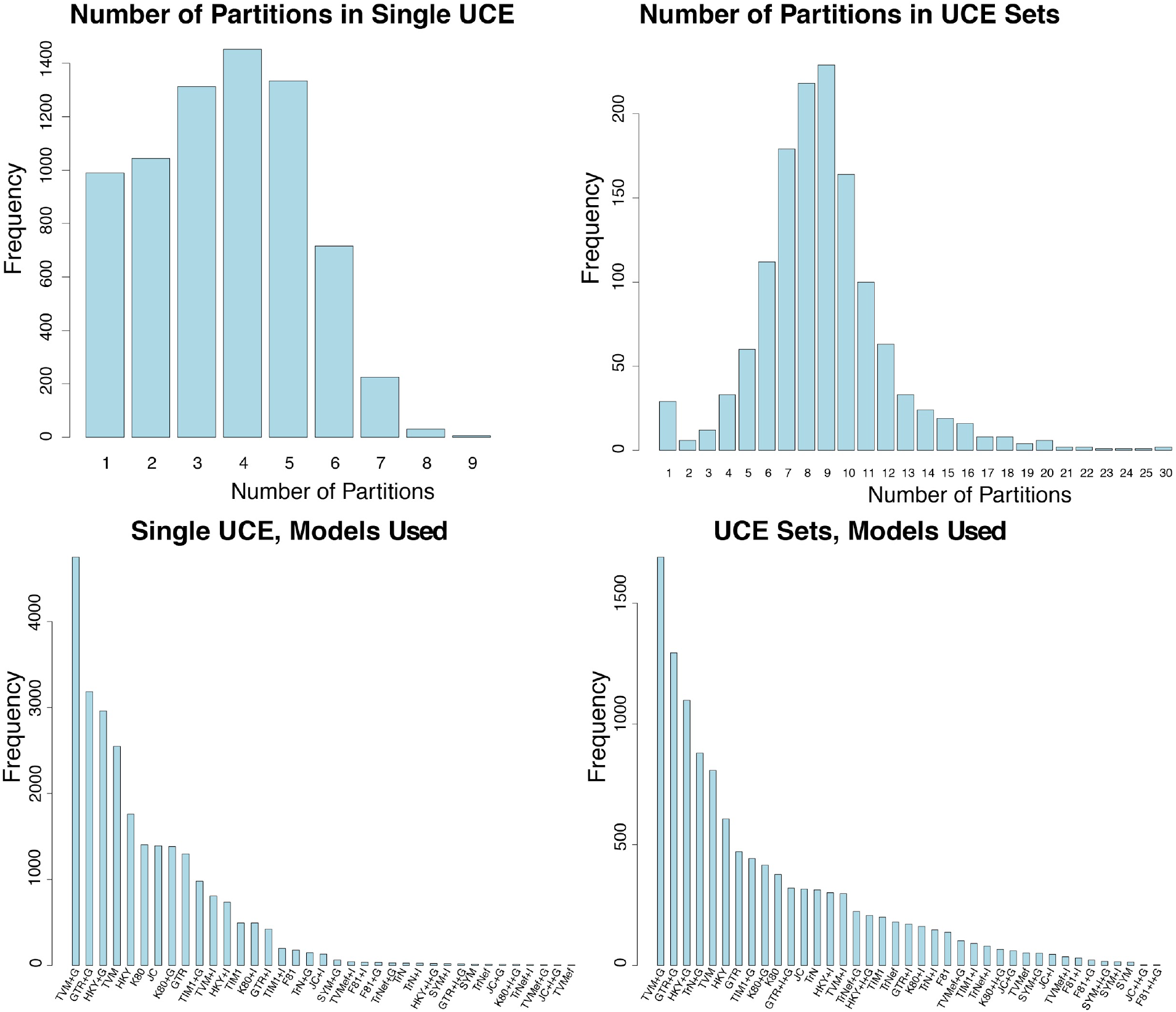
Barplot of UCE loci partitioning results. UCE “Sets” are concatenated UCEs which occupy a 25 kb non-overlapping sliding window bin, and “Singles” are those where only a single UCE occupies a 25 kb bin in the *P. sulphureomaculatus* base genome. Upper panels, barplot of the number of partitions found in a UCE locus. Lower panels, barplot of the number of times a particular model of nucleotide substitution was used in a partition.

### Divergence dating and Biogeography

The results of the MCMCTREE analysis estimated the root age from the 95% highest posterior density (HPD) of *Pachyrhynchus* to be between 25.1–17.9 Ma. Most *Pachyrhynchus* species have diversified within the last 5 Ma, and many species’ HPDs straddle the Pliocene-Pleistocene boundary (Fig. 4). The complete results of the MCMCTREE analyses are shown in Supplementary Fig. S4.

The results of the BioGeoBEARS analyses are shown in Table 3 and Fig. 4. The BioGeoBEARS analyses gave the greatest AICc model weight to the DEC+J model (Table 3). All models favored adding the founder event speciation parameter to the model, indicating that this mode of dispersal was significant in forming the broad scale biogeographic pattern of the genus. The root node of *Pachyrhynchus* was reconstructed as a joint range between Mindanao and Luzon. However, the descendant nodes were reconstructed as most likely Luzon. The subsequent radiations of lineages on Mindanao all descended from Luzon lineages. These separate Mindanao lineages represent five independent colonization events. The lineages of Mindoro, however, represent a mix of biogeographic lineages, three from Luzon and two from Mindanao. The single taxon from Panay represents a founder event from Luzon. Marinduque, although part of the Luzon PAIC, had two separate founder events one from Luzon and one from Mindoro. There was no evidence of back-colonization.

### Ancestral State Reconstruction

The equal rates (ER) model was selected as the one that best fit our data given the AIC weights (Table 4). The results of the ancestral state reconstructions of color pattern show that few clades were restricted to a single color pattern. The root of the tree was reconstructed as the spotted color pattern. The *“erichsoni”* and *“congestus”* clades, approximately one half of the taxa, were predominantly reconstructed as spotted. The changes from a spotted pattern to another state occurred along the terminal branches in this clade (Fig. 5). The other half of the taxa showed changes at deeper nodes, and their probabilities were not overwhelming given to one character state. One of the most striking observations from the reconstruction was that patterns, such as the net-like/Moroccan-tile pattern of *P. reticulatus,* occur independently in several different places in the tree. The unique rainbow color pattern (blues and yellows) also occurred in multiple clades (Figs. 2, 5).

**Figure 5.**
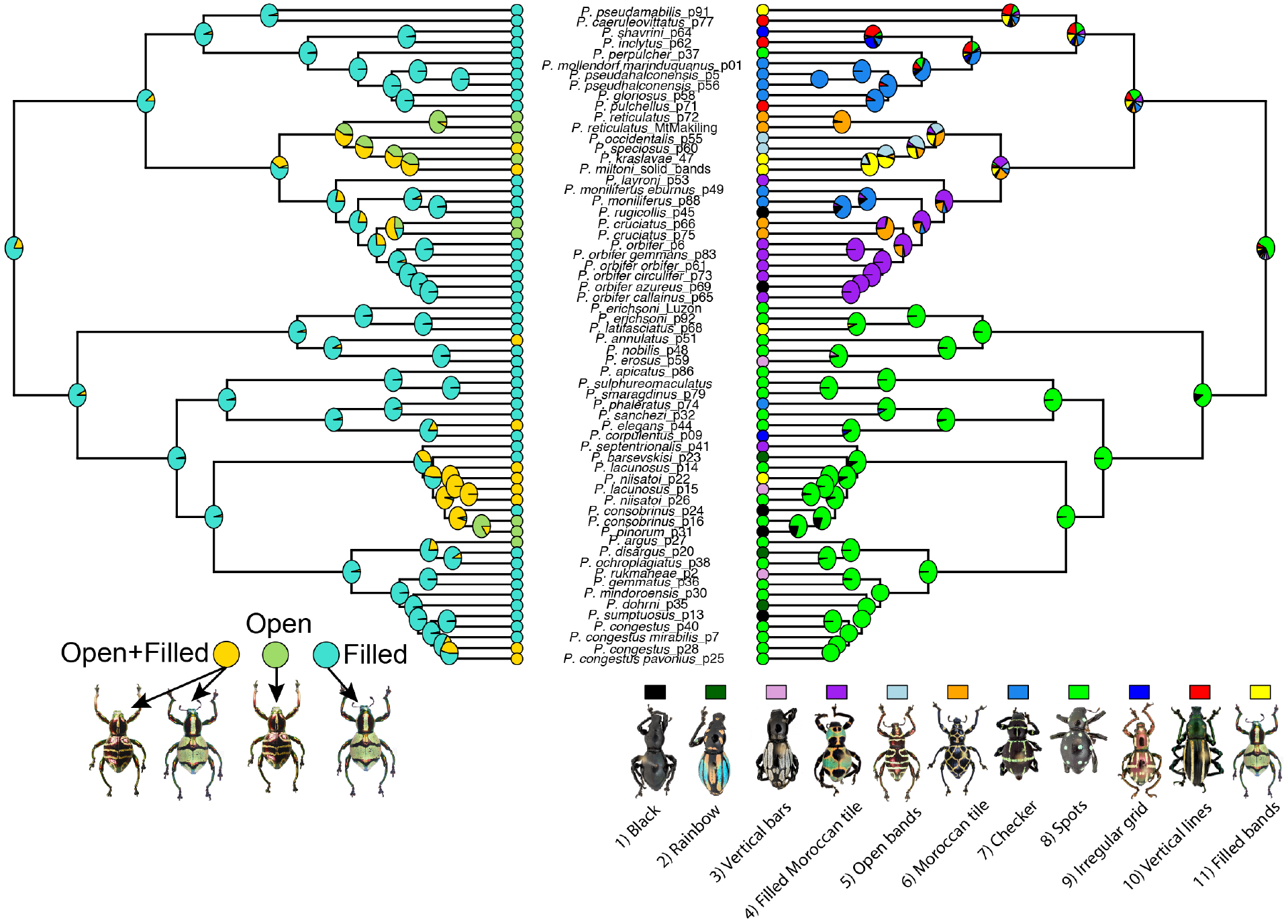
Ancestral state reconstruction of color patterns. **Left**, ancestral state reconstruction of polymorphic species states using *phytools “fitpolyMk”* function with the *“transient”* model. The character states were coded as “Open”, “Filled”, and “Open+Filled” where both states occur in a species. The polymorphic state can be associated with different color pattern states depending on the species (see lower right), for example “filled or open spots”, but never two different patterns within the same species e.g. “filled *bands* and open *spots”.* **Righ**t, ancestral state reconstruction using the *“fitMk”* function using the *“ER”* equal rates model. The 11 major color patterns observed in *Pachyrhynchus* are denoted in the lower right, colors correspond to those in the tree.

**Table 4.**
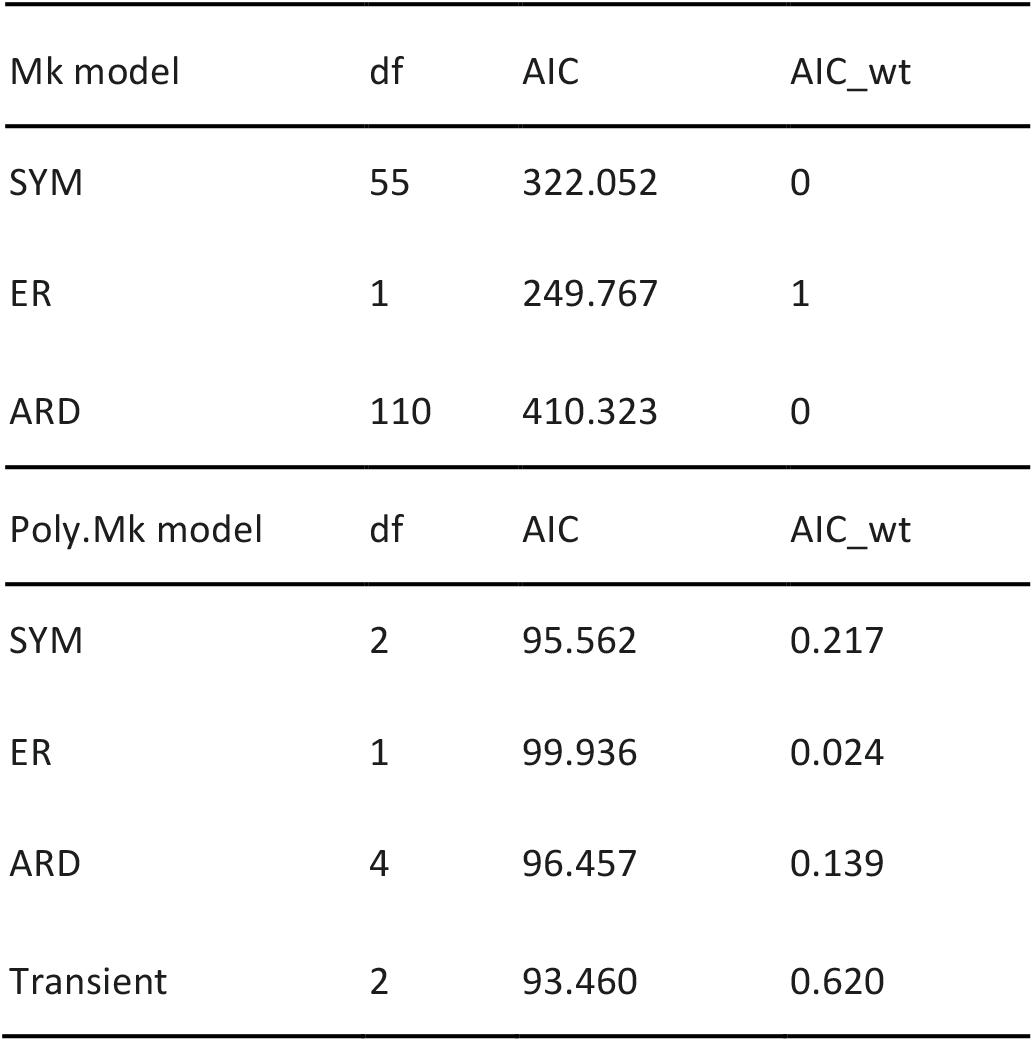
Color pattern model selection results.

| Mk model | df | AIC | AIC_wt |
| --- | --- | --- | --- |
| SYM | 55 | 322.052 | 0 |
| ER | 1 | 249.767 | 1 |
| ARD | 110 | 410.323 | 0 |
| Poly.Mk model | df | AIC | AIC_wt |
| SYM | 2 | 95.562 | 0.217 |
| ER | 1 | 99.936 | 0.024 |
| ARD | 4 | 96.457 | 0.139 |
| Transient | 2 | 93.460 | 0.620 |

For the analyses using the polymorphic Mk model framework, the *“transient”* model of transitions between different character states was selected as having the best fit to the data given the AIC weights. Here, the predominant pattern as well as the root node were reconstructed as having the filled bands character state (Fig. 5). There were six separate transitions from the filled state to the filled+open bands character state, two of these occurring along terminal branches. Polymorphic species occur in all major clades (Figs. 1–2, 5), except in the *“orbifer”* and *“gloriosus”* clades (Figs. 1, 5). In the *“orbiter”* clade, there was a transition from the polymorphic state to the open state at the node leading to *P. cruciatus.*

## DISCUSSION

We present the first phylogeny of the genus *Pachyrhynchus* based on 71 pinned and 16 ethanol preserved specimens using 10,108 UCE loci. Our results from both the concatenated and species tree analyses are highly supported with largely concordant topologies, demonstrating the benefit of Next Generation Sequencing and the value of historical museum specimens. With the phylogeny, biogeographic analysis, and ancestral state reconstructions of color patterns, we addressed how the wide array of phenotypes originated and broad scale evolutionary trends in this Müllerian mimicry system.

### Müllerian Mimicry in Pachyrhynchus

For this study, we functionally define Müllerian mimetic groups as two or more species of armored^23,24^ *Pachyrhynchus* weevils that share similar color patterns and are sympatric but do not share a most recent common ancestor. From our dataset, we identified 11 color patterns and at least nine distinctive mimetic groups in this system (Fig. 6). Members of each mimetic group closely resemble each other in their elytral patterns, and they also occur in sympatry, presumably sharing the same populations of predators (birds, lizards and frogs^23,25^). We found that most similar color patterns arose independently in distantly related taxa (Fig. 6), with a divergence time up to 20 MY between these taxa. For example, although *P. moniliferus* and *P. phaleratus* look superficially similar, they are not each other’s closest relatives, and are in fact at other ends of the tree (Fig. 6, top row, circle and square). As well, some color patterns do share common origins. For example, the *P. absurdus* species group and *P. speciosus* species group, co-occur throughout the greater Mindanao PAIC (Fig. 6, sixth row). This indicates that Müllerian mimetic evolution in *Pachyrhynchus* is mostly driven by convergence in conjunction with rare examples of shared ancestry. The mixed evolution of patterns has also been observed in *Heliconius* butterflies^12,86^.

**Figure 6.**
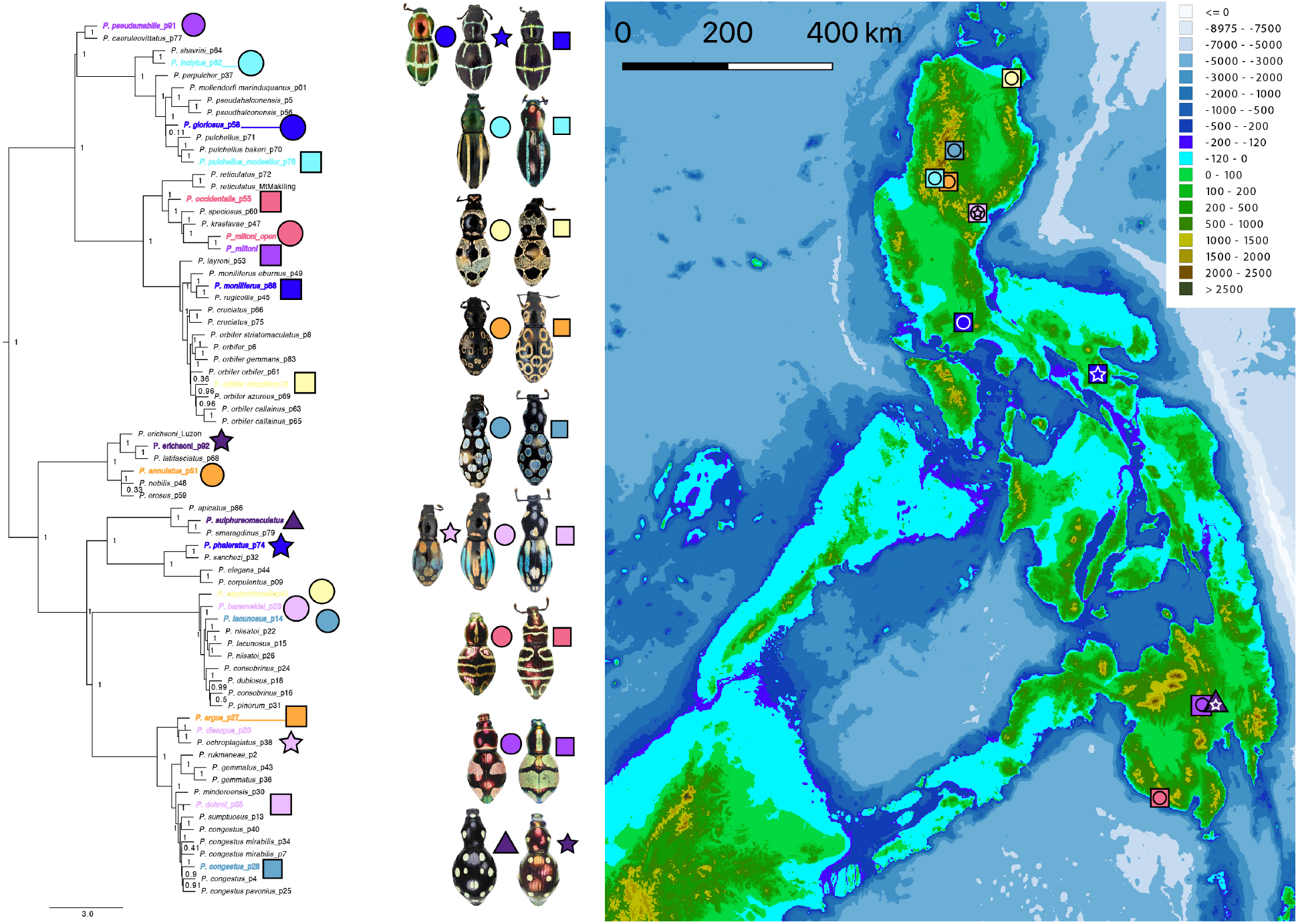
ASTRAL species tree of *Pachyrhynchus* with sympatric mimetic species. Colored symbols on phylogeny correspond to those in the central column and map. Node labels correspond to the LPP. **Central column**, mimetic species of *Pachyrhynchus.* **Right**, map of the Philippines, turquoise blue color denotes the −120m isobath, indicating land connection during Pleistocene glacial cycles.

While some members of the mimetic groups strongly resemble each other, others have only minor differences in pattern and/or color that are detectable by human eyes. Recent studies suggest that Müllerian mimicry might not be as clear-cut as originally proposed, and imperfect mimics and polymorphisms exist and may be common in nature^14,87–90^. These studies suggest that varied forms of mimics are able to persist due to limited cognitive capabilities of predators^91–93^ and/or predator avoidance of imperfect mimics due to the high cost of error^89,92,94^. Imperfect Müllerian mimicry adds additional complexity to the diversification process and phenotypic variation in *Pachyrhynchus.* For example, sympatric species in southern Mindanao, such as *P. erichsoni, P. miltoni,* and *P. pseudamabilis* all have a metallic red coloration to their cuticle, but differ in their maculations, spots or filled bands. (Fig. 6, bottom two rows).

### Biogeography of the Pleistocene Aggregate Island Complex (PAIC)

We find that all of the deeper lineages of the genus originated well before the PAIC was formed (Fig. 4). Additionally, within the major clades (Figs. 1–2), we find that most of their diversification events occurred during the Pliocene with some at the beginning of the Pleistocene (Fig. 4). This is significant because late Pliocene and early Pleistocene periods consisted of glacial/interglacial cycles^95–97^. These interglacial time periods should have promoted isolation between populations of *Pachyrhynchus.* Another period of higher sea level is the mid-Pliocene when it was 2-3°C hotter than today^98^, the interglacial periods would have caused significant isolation and perhaps promoted separate insular color morphs that would not have been compatible with neighboring larger PAIC islands when they were connected during lower sea levels. We see highly different color patterns today on such islands as Marinduque once connected to Luzon during the Pleistocene. The colonization history of such nearshore islands also contributes to differences from the larger PAIC islands. The divergence date for many species largely coincides with the mid-Pliocene time period, and as many *Pachyrhynchus* species are confined to higher elevations this may have influenced divergence on the larger PAIC islands (e.g., Luzon and Mindanao) as well. However, as there is a wide 95 HPD around these divergence times, we cannot entirely rule out early Pleistocene cycles as well. One striking pattern about the biogeography of *Pachyrhynchus* is that there is no back colonization of Luzon given its relative proximity and size to other major PAIC island groups. One factor that may have promoted this is that species colonizing an already inhabited landscape with other *Pachyrhynchus* species would not have had color patterns that matched the local fauna. However, this also could be a random effect, or an artifact of our sampling. Lastly, more precise geological data for when different islands emerged above sea level would greatly benefit Philippine biogeography.

### Insight into Pachyrhynchus Phylogeny and the Evolution of Color Patterns

Our phylogeny suggests that solely focusing on external morphology and color pattern may have driven inaccurate conclusions in previous taxonomic literature. For instance, *P. reticulatus* and *P. cruciatus* species share the superficial similarity of color pattern, while our phylogeny shows, with strong support, that they are distantly related, indicating convergence in color patterns. Although outside of the scope of this paper, our phylogeny could provide a great resource for taxonomic revision and species delimitation.

In addition to convergence, discrete polymorphism of patterns occurs frequently in populations and can complicate species identification if descriptions are based solely on morphology. Our phylogenetic results indicate that individuals with varied patterns should not be treated as separate species. For example, *P. miltoni* individuals may look different based on color pattern but are nearly identical genetically. We suggest that genetic data should be considered into future taxonomy and systematics of Pachyrhynchini beetles that have such complex mimetic evolution and biogeographic history. By integrating phylogenetic information to the taxonomy and systematics, the risk of creating synonyms or erroneous taxa will be diminished.

The type of binary polyphenism mentioned above is unusual in mimetic systems, with few examples in Coleoptera. One similar example occurs in the *Harmonia* ladybird beetles, but unlike *Pachyrhynchus,* the patterning is more of a continuum^99^. Additionally, in the ladybird beetle system, the color patterns are derived from pigments, and in *Pachyrhynchus* they are structural colors^27^. In Lepidoptera, discrete color polymorphism is more widespread and is known to occur in *Arctia plantaginis* tiger moths^100^, between the sexes of *Neophasia terlooii* butterflies, males are white and females are orange^101^, as well as in *Heliconius*^12^.

We suggest that selection of color patterns is frequency-dependent. This hypothesis is supported by casual observations of *Pachyrhynchus* specimens in the CASENT collection, we counted the relative abundances of beetles with particular patterns in some well-studied localities. For instance, *P. moniliferus* is a widespread, relatively common species found in southern Luzon. It co-occurs with several other species with more restricted ranges. For example, in Iriga, Camarines, Luzon, 69 *P. moniliferus* specimens (CASENT) were collected, but only three of the larger *P. phaleratus* (Fig. 6 top row, star and square) were collected at the same location and time period (May–Oct. 1931). In addition, at Mt. Makiling, Luzon, (May–Dec. 1930-31), 38 specimens of *P. gloriosus* were collected compared to 225 *P. moniliferus* (Fig. 6 top row, circle and square). Because all *Pachyrhynchus* species have a hard cuticle (often bending pins during preparation)^23^, but have many different color patterns, frequency dependent selection is likely acting on the color pattern and not cuticular hardness. In the case of polymorphic species, frequency dependent selection may also be a cause of the observed frequencies. For example, in *P. miltoni,* found in the southern Davao City Province of Mindanao, the ratio of the filled band morphs to the open band morphs is 35:2, but in neighboring populations near Mt. Apo, the open-banded morph is most abundant (CRC and CASENT collections). Although more methodical studies should be done to control for artifacts caused by uneven sampling, different years of collection, seasonality, etc.; including information about polymorphism helps to elucidate how particular patterns evolved. For instance, *P. reticulatus* species with open bands were descended from polymorphic populations of *P. reticulatus* inferred from our ancestral state reconstructions (Fig. 6). Because these beetles are flightless with patchy distributions, (some only restricted to a single mountain), the relative frequencies of these color morphs are likely to be fixed in isolated populations and is a possible cause for speciation. This could perhaps explain why we find convergent color patterns between allopatric species that are not each other’s closest relatives.

### UCE Partitioning

We have demonstrated that the universal use of a single model of nucleotide substitution is inadequate to accommodate site rate heterogeneity across UCE sets. The preferred use of multiple modes within a UCE locus has also been observed when constructing gene trees with MrBayes^62^. Additionally, we found that using more than three partitions was selected as optimal. While UCE loci have been treated as varying symmetrically around the central core^62,63,102^, we find that often this is not the case as a variable, even number of partitions was selected. This suggests that flanking regions tend to be highly variable and that treating them as symmetrical units is a suboptimal partitioning design. More analyses are required to compare the design of Tagliacollo and Lanfear 2018^63^ to the design we propose here to make a decisive decision of which scheme is optimal for UCE partitioning. The partition method we employ is the only one that accommodates the combination of neighboring or co-genic UCEs^51^ (the method of Tagliacollo and Lanfear 2018^63^ could be easily updated). The asymmetry in variation away from the UCE core is perhaps due to the way in which UCE loci are treated in the alignment process, mainly that difficult to align regions are trimmed in a less symmetric manner around the core of the locus.

### Future Directions

Several unique features add intriguing complexity to our study system and, thus, inspire further studies. Color patterns in *Pachyrhynchus* are formed by the arrangement of scales, and the different scales’ colors result from light reflectance on photonic crystals coupled with the background color of the elytra. The colors are structural^27^, differing from the pigment-based colors of most other Coleoptera^14^ or *Heliconius* butterflies. This indicates a different genetic pathway that underlies their evolution and diversification. Our phylogeny provides a robust basis for further research to uncover the genetic mechanism controlling structural coloration in an evolutionary framework.

The armored exoskeleton is essential for a weevil’s survival. Thick cuticle formation relies on the generation of its precursor amino acid, tyrosine, produced by obligate bacterial endosymbionts of the *Nardonella* lineage^26^. *Nardonella* has a much-reduced genome of 0.2 Mb and lacks genes responsible for most metabolic pathways; they rely on the beetle’s metabolic output for survival^26^. Interestingly, *Pachyrhynchus’* elytra and cuticle are initially soft and easily deformed when they are teneral (soft bodied) adults, but the color patterns are apparent as soon as weevils emerge from the pupae^26^ (personal obs. A Cabras). Their bold patterning likely provides protection from predators that have learned to avoid the aposematic signal in older adults. Predator avoidance during vulnerable stages is likely essential for survival until reproduction. More ecological studies on population densities of sympatric *Pachyrhynchus* species and predator preferences can also help to elucidate color pattern evolution on a landscape scale.

In summation, we clearly demonstrated that many of the mimetic color patterns observed in sympatry are due to convergent evolution and not simply due to inheritance. Based on our observations in natural history collections, we hypothesize that convergence between these independent color pattern forms is likely driven by frequency dependent selection. Lastly, the use of a UCE design specific to the tribe Pachyrhynchini was highly successful in resolving the relationships between our taxon.

Our study presents an interesting system of Müllerian mimicry and provides a framework for an integrative approach to study other similar systems.

## Acknowledgments

This research was funded in part by a generous gift from to the C.A.S. from Will and Margaret Hearst for supporting the 2011 Filipino-American Hearst Biodiversity expedition to Luzon. We were also funded in part through NSF:DEB award number 1856402 made to MHVD. We would like to thank the Ruth Tawan-tawan, Ceso II of the Philippines’ Department of Environment and Natural Resources Region XI for help with the Gratuitous and export permits. We would also like to thank the University of Mindanao for the mobility support, and Milton N. Medina and Chrestine Torrejos of U.M. for help collecting specimens. We would also like to thank Sarah Crews and Alejandra Hernandez-Agreda of C.A.S. for help with the manuscript text. We would like to thank Jim Henderson C.A.S. for help with the 10X assembly.

## Supplemental Figures

**Supplementary Figure 1.**
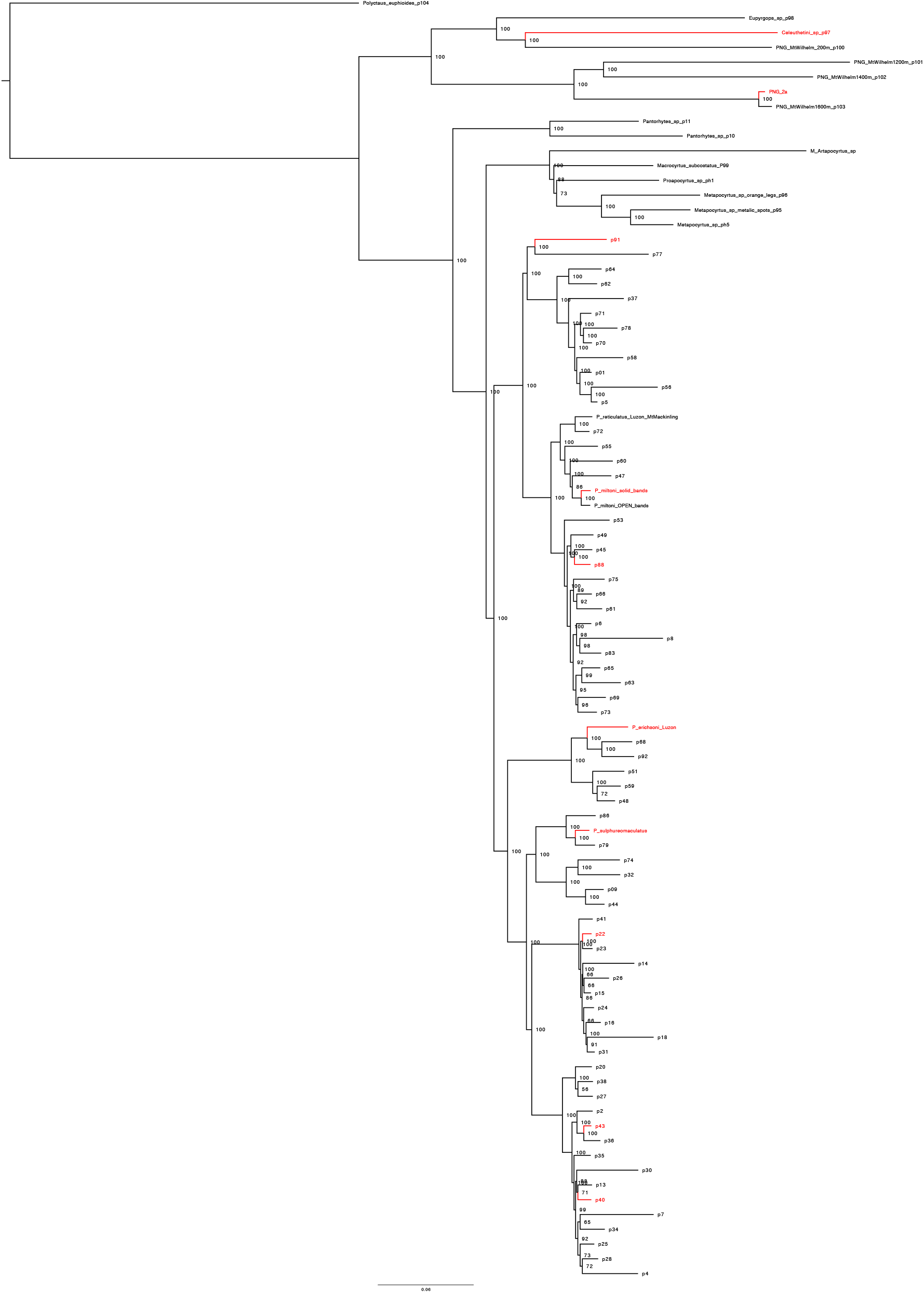
*Pachyrhynchus* concatenated ML phylogeny constructed with RAxML-NG. Node labels are bootstrap support values. Branches in red are the taxa used in probe design.

**Supplementary Figure 2.**
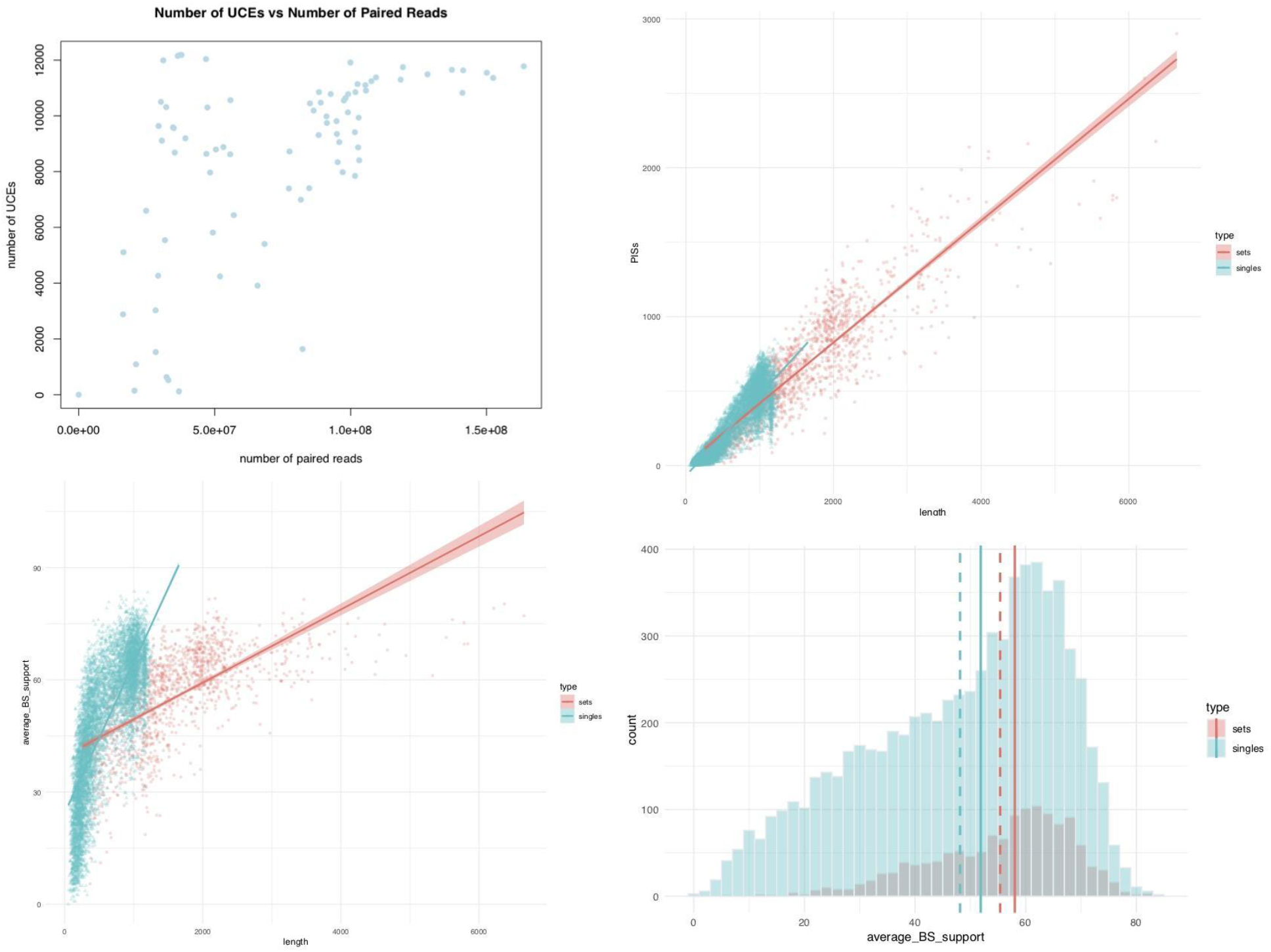
**Upper left** panel, Number of UCEs recovered vs number of paired reads. For the UCE type, the ‘‘sets” are concatenated UCEs found in a 25kb non-overlapping sliding window bin, and UCE ‘‘singles” are those where only a single UCE occupies a 25kb bin in the *P. sulphureomaculatus* base genome. **Upper right**, Phylogenetically informative sites vs length of UCE locus. **Lower left**, Average bootstrap support per locus by UCE type vs length of locus. **Lower right**, Average bootstrap support per locus by UCE type. Solid vertical lines are the mean and dashed vertical lines are the median by UCE type.

**Supplementary Figure 3.**
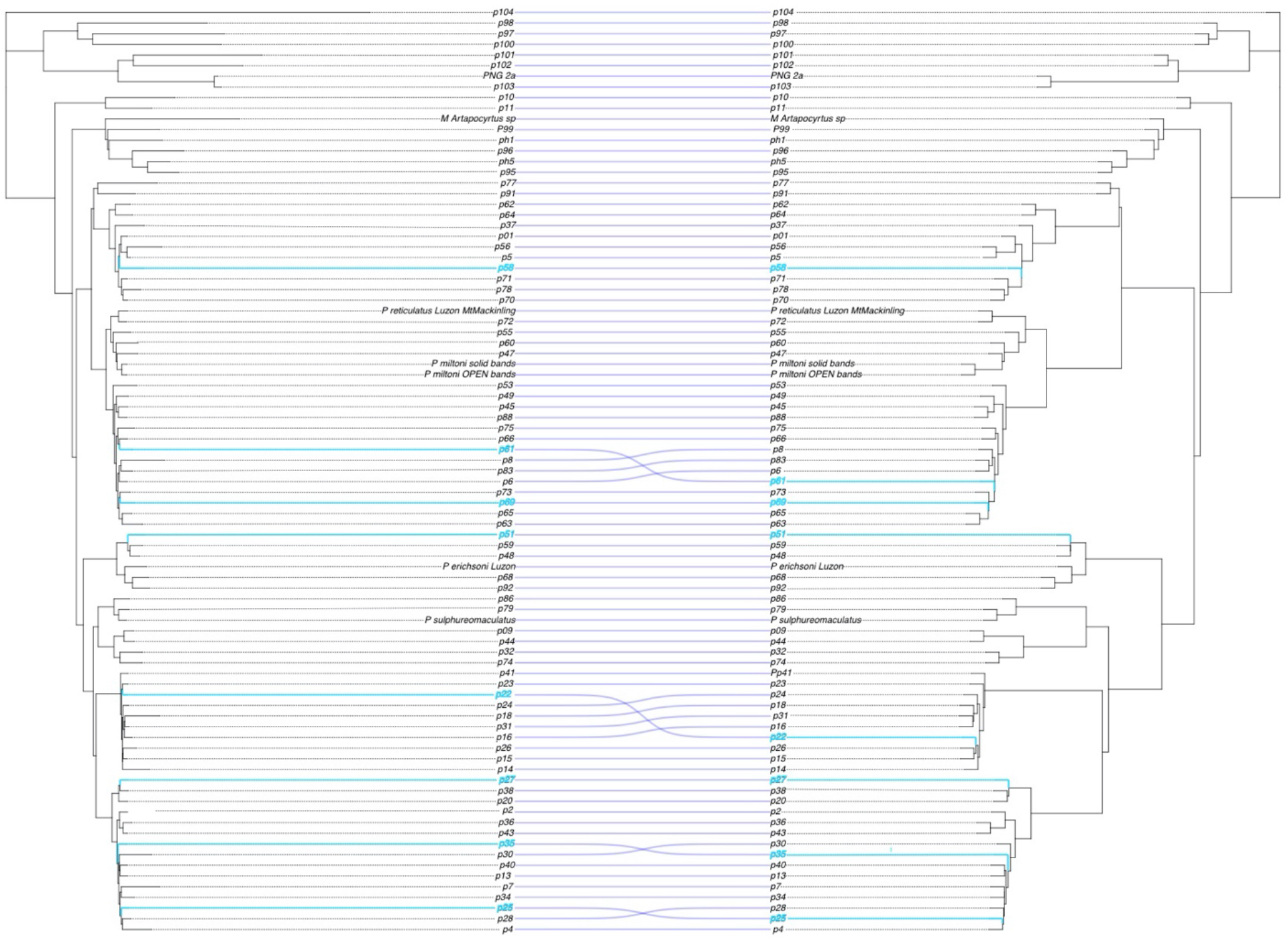
RAxML concatenated phylogeny on left, ASTRAL species tree on right.

**Supplementary Figure 4.**
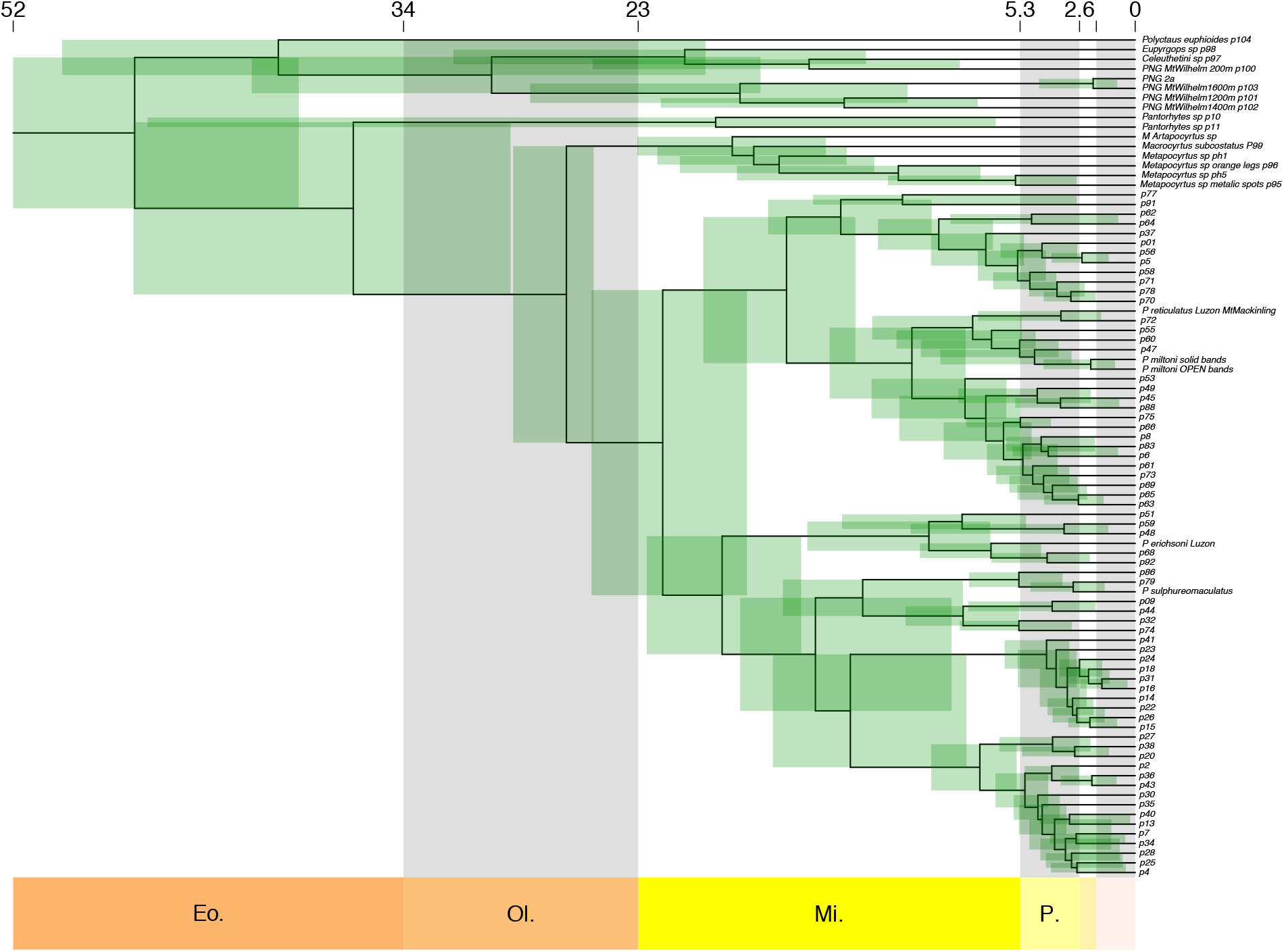
*Pachyrhynchus* chronogram constructed using MCMCTREE. **Nucleotide Models Used:** GTR+I, GTR+G, F81, F81+G, SYM, F81+I, JC, HKY, K80, HKY+I+G, K80+I, SYM+G, SYM+I, K80+G, GTR, HKY+G, SYM+I+G, HKY+I, F81+I+G, JC+G, GTR+I+G, JC+I, K80+I+G, JC+I+G, TIM, TVM, TVMef, TrN, TrNef, TIM+G, TVM+G, TVMef+G, TrN+G, TrNef+G, TIM+I, TVM+I, TVMef+I, TrN+I, TrNef+I, TIM1+I, TIM1,TIM1+G

